# SARS-CoV-2 variants of concern Alpha, Beta, Gamma and Delta have extended ACE2 receptor host-ranges

**DOI:** 10.1101/2021.11.23.469663

**Authors:** Nazia Thakur, Giulia Gallo, Joseph Newman, Thomas P. Peacock, Luca Biasetti, Catherine N. Hall, Edward Wright, Wendy Barclay, Dalan Bailey

**Affiliations:** The Pirbright Institute, Guildford, Surrey, GU24 0NF, United Kingdom; Nuffield Department of Medicine, The Jenner Institute, Oxford, OX3 7DQ, United Kingdom; Department of Infectious Disease, Imperial College – London, United Kingdom, W2 1PG; School of Psychology and Neuroscience, University of Sussex, Falmer, BN1 9QH, United Kingdom; Viral Pseudotype Unit, School of Life Sciences, University of Sussex, Falmer, BN1 9QG, United Kingdom

## Abstract

Following the emergence of SARS-CoV-2 in China in late 2019 a number of variants have emerged, with two of these – Alpha and Delta – subsequently growing to global prevalence. One characteristic of these variants are changes within the Spike protein, in particular the receptor binding domain (RBD). From a public health perspective these changes have important implications for increased transmissibility and immune escape; however, their presence could also modify the intrinsic host-range of the virus. Using viral pseudotyping we examined whether the variants of concern (VOCs) Alpha, Beta, Gamma and Delta have differing host ACE2 receptor usage patterns, focusing on a range of relevant mammalian ACE2 proteins. All four VOCs were able to overcome a previous restriction for mouse ACE2, with demonstrable differences also seen for individual VOCs with rat, ferret or civet ACE2 receptors, changes which we subsequently attribute to N501Y and E484K substitutions within the Spike RBD.

## Main text

SARS-CoV-2, the β-coronavirus responsible for the Covid-19 pandemic, is thought to have emerged from a bat reservoir, potentially via an as yet unidentified intermediate mammalian host. These conclusions are supported by knowledge of the origins of SARS-CoV-1 as well as the sequence data of related viruses isolated in bats e.g., the RaTG13 isolate [1]. Recently, Temmam et al., identified sarbecoviruses in bats that are nearly identical within the Spike RBD, which adds further weight to this conclusion and highlights that coronaviruses with high human ACE2 affinity are actively circulating in wild reservoir populations [2]. Interestingly, experimental infection of ferrets, mice, bats, primates and other animals, together with natural infections in a range of species including cats, dogs and mink indicate these viruses may have a broader host-range than bats and humans [3–6]. Understanding SARS-CoV-2 infection in animals is important for three main reasons. Firstly, to assess the direct risk to livestock, companion animals and wildlife. Secondly, to examine whether these animals can act as secondary reservoirs for SARS-CoV-2. And lastly, to help establish and validate animal models for Covid-19 which can then be used for the development of therapeutics and vaccines, and to better understand the mechanisms of viral pathogenesis. To support these research endeavours, we and others have shown that SARS-CoV-2 Spike has a broad tropism for mammalian ACE2 proteins [7–9]. Using lentiviral pseudotyping combined with live virus experiments we showed that SARS-CoV-2 can use a wide range of mammalian ACE2s including dog, cat, cattle, sheep, pangolin and rabbit, but is restricted with rat, ferret and a subset of bat and bird receptors [7, 10].

Importantly, the continued evolution of SARS-CoV-2 in human populations has led to the emergence of variants, a natural result of RNA virus replication. Informed by epidemiological, virological and immunological data, public health bodies such as the World Health Organisation (WHO) and Public Health England (PHE) have assigned some as variants of concern (VOCs). Over the course of the pandemic two of these VOCs (B.1.1.7 [Alpha] and B.1.617.2 [Delta]) have independently risen to prominence, rapidly replacing the previously circulating strain across multiple regions (G614) [11]. The mechanisms underpinning this replacement appear to correlate with continued evolution to the human host, through increases in particle infectivity, replicative capacity, transmission potential, innate immune antagonistic properties and potentially immune escape [12–16]. However, whether these variants have expanded host-ranges and the consequences for the ongoing pandemic, in terms of the factors described above, are not well characterised.

Using a lentiviral (HIV-1) pseudotyping approach for SARS-CoV-2 (described previously, [7, 17]) we investigated whether the Spike from four different VOCs (Alpha [Spike mutation profile: L18F, Δ69-70, Δ144, N501Y, A570D, D614G, P681H, T716I, S928A, D1118H], Beta [L18F, D80A, D215G, Δ242-244, K417N, E484K, N501Y, D614G, A701V], Gamma [L18F, T20N, P26S, D138Y, R190S, K417T, E484K, N501Y, D614G, H655Y, T1027I, V1176F] and Delta [T19R, G142D, Δ156-157, R158G, L452R, T478K, D614G, P681R, D950N]; Figure 1 A-D; left panel) had altered ACE2 receptor usage across multiple hosts, when compared to a sequence derived earlier in the pandemic (WT/ D614/ Wuhan). Expression constructs for the various Spike proteins were constructed in a background lacking the final 19 amino acids of the C-terminal cytoplasmic tail (Δ19; equivalent to K1255*STOP) to enhance spike incorporation and facilitate increased pseudotyping efficiency, either by site directed mutagenesis or gene synthesis [18]. In addition to human ACE2, other host ACE2s were chosen based on their relevance to previous sarbecovirus emergence (civet), their continued use as animal models for SARS-CoV-2 pathogenesis and transmission studies (ferret, hamster, mouse), as well as a perceived potential to act as a reservoir for human-derived viruses (rat [through excreted virus in sewage], pig [established reservoir for Nipah and influenza]). Briefly, pseudotyped virus for WT D614 and the four VOCs was generated in HEK293s and used to infect BHK-21s over-expressing the various ACE2s. Luciferase activity, a quantitative read-out proportional to pseudo-viral entry, was then assayed at 2 days post-infection. These values were then normalised based on the results of product-enhanced reverse transcriptase (PERT) assays for each pseudovirus preparation, a method used to standardise results when comparing different preparations of lentiviral pseudotypes. Experiments were repeated three times and the data for each VOC collated and compared to WT D614 (Figure 1). For human ACE2, the only statistically significant difference (t-test) in entry was observed for the Beta VOC, which showed a small increase when compared to WT D614 (Figure 1B). Usage of mouse ACE2 by all four VOCs was significantly increased when compared to WT D614 (Figure 1 A-D), in keeping with previous findings that early SARS-CoV-2 isolates were restricted with this specific receptor [7]. Of note, the increase in mouse ACE2 usage was less notable for Delta, the only non-N501Y-containing variant in our study. Similarly, the SARS-CoV-2 Spike – rat ACE2 restriction we reported previously [7] was only overcome by N501Y-containing VOC Spikes (Alpha, Beta, Gamma), with no significant increase seen with Delta (Figure 1). For civet ACE2 significant increases in entry were observed only with the Beta VOC, while for ferret ACE2 Beta and Gamma were both significantly higher than WT D614. Interestingly, no significant differences were observed when hamster or pig ACE2 were used as receptors although in our 2020 study neither of these receptors were restrictive to early isolates of SARS-CoV-2 entry, unlike rat, mouse, civet and ferret ACE2 [7]. Analysis of the same data on radar plots highlights the extended host-range of the N501Y-containing VOCs (Alpha, Beta and Gamma), while simultaneously demonstrating the similar host-range profiles of WT D614 and Delta (Figure 1 A-D right panels).

**Figure 1:**
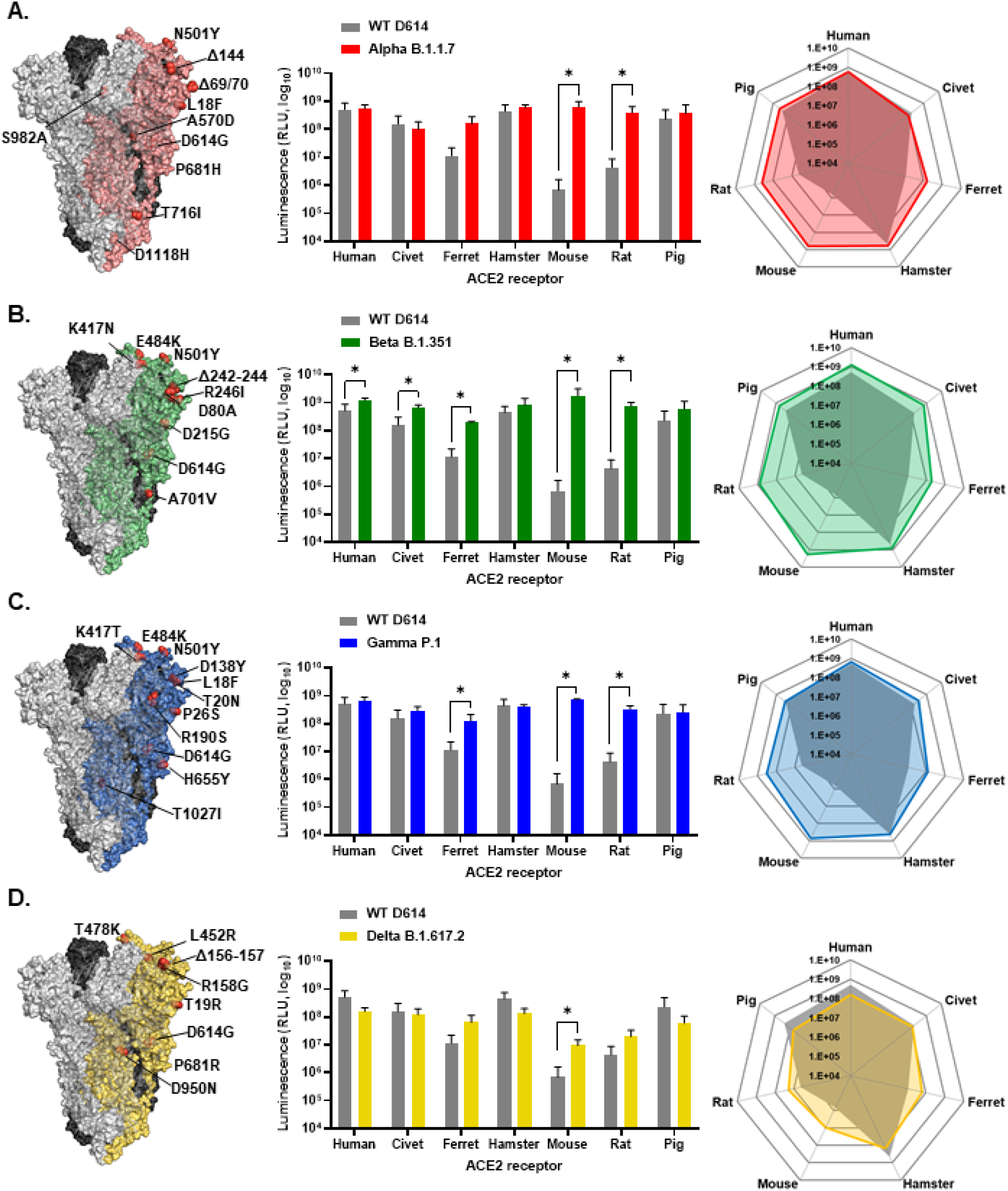
ACE2 receptor usage screening of SARS-CoV-2 variants of concern (VOC) Spike proteins. Structural maps and receptor usage analysis are shown for SARS-CoV-2 VOCs (A) Alpha (B.1.1.7), (B) Beta (B.1.351), (C) Gamma (P.1) and (D) Delta (B.1.617.2), compared to WT D614 (Wuhan). **Left panels:** Amino acid differences between the VOC and WT D614 were highlighted on a trimeric surface structure of SARS-CoV-2 Spike using Pymol (PDB: 6XM4). Spike monomers are coloured according to the indicated VOC; red (Alpha), green (Beta), blue (Gamma), yellow (Delta), with substitutions identified on the coloured structure only. **Middle panels:** Receptor usage was screened using HIV-1 lentiviral pseudotypes bearing the indicated Spike proteins with HEK293 target cells expressing the indicated ACE2 protein. Viral entry was measured by assaying luciferase activity (RLU) using the BrightGlo reagent (Promega). **Right panels:** The same data was replotted on radar plots to illustrate the broadening host-range of the Alpha (red), Beta (green) and Gamma (blue) VOCs, as opposed to Delta (yellow). In each plot the signal for WT D614 is provided in grey (transparent polygon). For entry assays, BHK-21 cells were plated at 5×10^5^/well in a 6-well dish and transfected with the indicated ACE2 expression plasmids (human, pig, rat, hamster, ferret, civet, mouse) 24 hours later. The next day, cells were harvested using 2mM EDTA-PBS and diluted to 2×10^5^/ml with 100μL being seeded into white-bottomed 96-well plates and incubated for 24 hrs at 37oC, 5% CO2. The media was removed from plated cells and infected with 100μl of the indicated pseudotyped virus and incubated for 48 hours. Firefly luciferase was quantified on a GloMax Multi+ Detection System using Bright-Glo (Promega). Individual experiments were performed in biological triplicates. These experiments were then repeated on three separate occasions (on different days). PERT-standardised RLUs were normalised between experiments based on the average RLU calculated across the whole plate, with the mean RLU for each condition then calculated and plotted. To facilitate interpretation and comparison of the dataset the same WT D614 data has been plotted for each VOC. Statistical significance was measure by students t-test.

To examine the role of individual amino acid changes in Spike in overcoming host-range restrictions we subsequently focused on the mouse, rat and civet ACE2 interactions. Plasmids expressing SARS-CoV-2 Spike mutations found in various VOCs were constructed in a WT Δ19 background and used to generate pseudotyped viruses. Subsequent infection of mouse ACE2 expressing cells confirmed that a single N501Y change was sufficient to overcome the WT D614 restriction with this receptor, allowing entry equivalent to the Alpha VOC (Figure 2A). Changes to the furin cleavage site (P681H, found in Alpha) or deletions in the N-terminal domain (NTD; Δ69-70, Δ144), however, appeared to have little appreciable effect on entry, with similar results being observed for rat ACE2 (Figure 2B). Of note the Δ69-70 mutations were performed in a N501Y-containing background, which could potentially obscure synergistic effects between these two mutations. Interestingly for civet ACE2, the N501Y or K417N changes were inhibitory, with E484K, in contrast, increasing entry efficiency of the respective SARS-CoV-2 pseudotype (Figure 2C). The small, but repeatable, increase in civet ACE2 usage with the Beta VOC Spike (Figure 1B) may therefore be a combinatorial effect of these mutations, with E484K compensating for the inhibitory properties of the 501 and 417-specific changes. Accordingly, an Alpha + E484K Spike-based pseudotype was able to rescue the defect in civet ACE2 usage seen with Alpha alone (Figure 2C; left panel).

**Figure 2:**
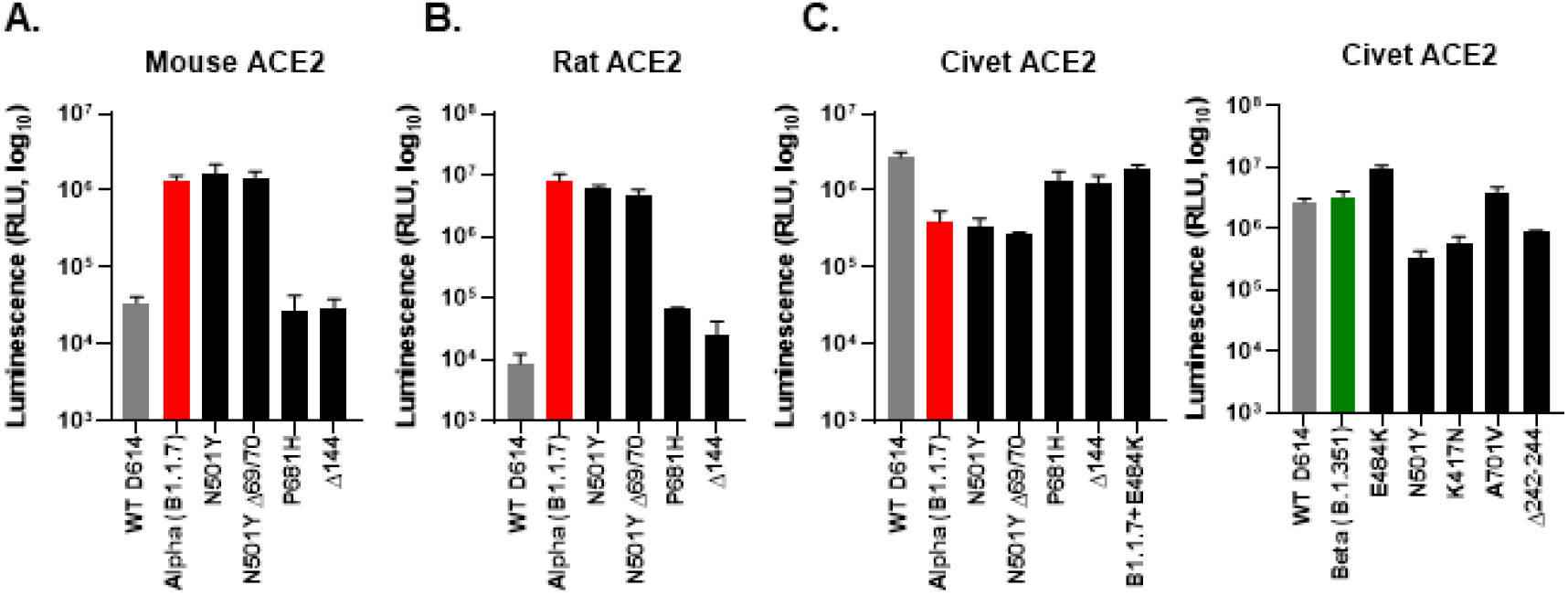
Mutational analysis of the RBD identifies N501Y as a critical amino acid for extending the host-range of SARS-CoV-2 Spike to rodents. The entry efficiency of pseudotypes bearing SARS-CoV-2 Spikes with individual or combinations of mutations found in the Alpha and Beta VOCs was assayed in mouse (A), rat (B) or civet (C) ACE2 expressing cells and compared to Alpha, Beta or WT D614. Viral entry was measured by assaying luciferase activity (RLU) using the BrightGlo reagent (Promega). Individual experiments were performed in biological triplicate, with the mean RLU for each condition then calculated and plotted.

The continuing evolution of SARS-CoV-2 in human populations raises significant concerns. Principally, these relate to the human pandemic and human disease, e.g., antigenic escape from vaccine- or natural infection-derived immunity, or the acquisition of enhanced transmission potential or increased pathogenicity. Separately, however, the question of whether SARS-CoV-2 will develop enhanced reverse zoonotic potential is also important. The emergence of both the Alpha and Delta VOCs provides clear evidence that a relatively small number of changes within the genome can have far reaching epidemiological consequences for the human pandemic. To our knowledge these VOCs have, however, not yet been linked to any significant increase in spill-over back into livestock, companion animals or wildlife; for which there may be various interpretations. This may reflect a lack of sampling, improved understanding and enhanced biosecurity at the human-animal interface, a consistently low frequency of spill-over or simply the absence of any dramatic change in disease phenotype following infection of animals with VOCs.

Regarding infection, our data show that from an ACE2 receptor usage perspective the host-range of Alpha, Beta and Gamma (and to a lesser extent Delta) is broader than the virus that initially emerged in China (WT D614). Several host ACE2 proteins that were previously refractory to viral entry (mouse, rat and ferret) are used more efficiently by VOC Spikes (Figure 1). This change in tropism is attributable to specific amino acid changes in the RBD of Spike, in particular N501Y (Figure 2). This is consistent with previous work showing amino acid substitutions within the RBD overcome restriction with murine ACE2. These mutations were identified by structure-led approaches [19] or following passage of SARS-CoV-2 directly in mice [20], results which have been subsequently confirmed *in vitro* [21, 22]. Although these changes are not identical, they fit within an overall pattern that small adaptations in Spike are needed to overcome specific incompatibilities between the viral attachment protein and non-cognate receptors. We observed a similar trend during adaptation of SARS-CoV-2 in ferrets, with Spike mutations Y453F, F486L and N501T leading to increased ferret ACE2 usage [10]. This mirrors the situation with natural infections in mink, a related mustelid, where a similar array of mutations has repeatedly arisen [23, 24] leading to severe restrictions on the farming of mink and the mass culling of animals. Importantly, these ferret or mink acquired mutations are unlikely to pose an increased threat to human populations as these changes either inhibit human ACE2 usage (Y453F) and/or have little antigenic impact [10].

Whether SARS-CoV-2 VOCs are more pathogenic in rodents is an area of ongoing investigation. Montagutelli et al., showed that the Beta and Gamma VOCs replicated to higher titres than a B.1 virus in the lungs of young adult mice [25] with Horspool et al., showing similar results in K18-human ACE2 transgenic mice with Alpha and Beta VOC infected animals showing higher cumulative clinical scores when compared to the first wave WA-1 strain [26]. Similar results were also reported by Shuai et al in mice and rats [27]. This has contributed to speculation that rodents could play an important role in the transmission of SARS-CoV-2 to people [28]; however, these data also have important ramifications beyond reverse zoonosis. For example, significant weight is given to animal model data on SARS-CoV-2 pathogenicity during antiviral screening or vaccine development [6, 29]. If, for example, N501Y containing VOCs cause increased pathogenicity because they are now able to use mouse ACE2 more efficiently, this could complicate or potentially invalidate comparisons between WT D614 and VOC-based mouse data and more broadly correlations with human data, where the difference in human ACE2 usage is marginal. In addition, those animal models that endogenously express restricted ACE2s (ferret, mouse and rat) are likely to drive in-animal adaptation, as we witnessed with ferrets [10], which further complicates conclusions on viral phenotypes such as transmission potential. Fortunately, in hamsters which have emerged as a highly tractable model for SARS-CoV-2, the situation appears less convoluted as differences in hamster and human ACE2 usage between WT D614 and VOC pseudotypes look comparable (Figure 1).

While in ferrets and mice the increased capacity for SARS-CoV-2 VOCs and other variants to replicate in these hosts correlates with changes in the Spike protein, it is difficult to make a broader set of conclusions. Why certain hosts develop disease and others don’t and how this could impact the Covid-19 pandemic is still a relatively unanswered question. A relevant example is pigs, which we have established encode a cognate receptor for SARS-CoV-2 entry (Figure 1). Pigs have proven time and again to be an adequate reservoir for human tropic viruses (e.g. influenza and Nipah), yet are apparently refractory to SARS-CoV-2 infection [30, 31]. Recently, we demonstrated tissue-specific differences in ACE2 expression between various animals, e.g. ACE2 was present in the nasal mucosa epithelium of Eptesicus serotinus (serotine bat) but not in pigs (*Sus scrofa domestica*) [32] which may provide some mechanistic insight into the varying susceptibility of hosts to SARS-CoV-2. Beyond entry, various species-specific restrictions, for example at the level of the innate immunity response, may also be playing a role.

Nevertheless, the fundamental question that remains is whether existing or emerging SARS-CoV-2 variants can establish themselves in an animal reservoir in a way that is consequential to management of the human pandemic. Aside from mink infections, where the virus likely had to adapt in order to use the mustelid receptor, other relevant examples include spill-over and onward transmission in free-living and captive white tailed deer [33] as well as emerging evidence of cryptic SARS-CoV-2 lineages in wastewater that may be linked to establishment of a rodent reservoir [34]. Whilst this evidence and that of human-companion animal transmission, e.g. for cats and dogs [35], seems strong the reverse scenario, i.e. infection of humans by SARS-CoV-2 infected animals appears much less common, albeit not totally absent [23, 24]. The conceivable ‘worst case scenario’ for SARS-CoV-2 reverse zoonosis is that the virus establishes itself in a new reservoir and at such a level that antibody selection pressure takes place and/or prolonged antigenic drift leading to escape mutants that are relevant to immune human populations. Our previous data indicated that SARS-CoV-2 was a ‘generalist’ with a broad host-range [7]. Significantly, the VOCs have even broader host ranges and, in the majority of cases, the amino acid changes involved e.g., N501Y, which enhances human ACE2 binding (providing evidence of ongoing human adaptation), do not lead to a loss of activity with other host ACE2s. In other words, the ‘generalist’ nature of SARS-CoV-2 is being maintained and extended over time. Whether this trend will continue and whether this increases the probability of human-relevant reverse zoonosis events remains to be determined.

## Acknowledgements

This work was supported by the G2P-UK National Virology Consortium funded by MRC funded grant G2P-UK; A National Virology Consortium to address phenotypic consequences of SARS-CoV-2 genomic variation (MR/W005611/1). DB, NT, JN and GG were also funded by The Pirbright Institute’s BBSRC institute strategic programme grant (BBS/E/I/00007038 and BBS/E/I/00007034) with NT receiving studentship support from BB/T008784/1. EW, CH and LB were supported by the MRC grant MR/V036750/1.

